# Internalization-Responsive Nano-PLGA For Image-Guided Photodynamic Therapy Against HER2-Positive Breast Cancer

**DOI:** 10.1101/2023.02.21.529435

**Authors:** Anna S. Sogomonyan, Sergey M. Deyev, Victoria O. Shipunova

## Abstract

Oncotheranostic nanoagents are powerful artificial tools that significantly outperform traditional antitumor therapies. Among the wide variety of nanoagents, polymer nanoparticles have proven to be one of the most successful candidates for translation into clinical practice. Here we developed an oncotheranostic platform that is based on poly(lactic-co-glycolic acid) nanoparticles and possesses red/green light dual-activated diagnostic/therapeutic properties for targeted photodynamic therapy of HER2-positive aggressive breast cancer. PLGA nanoparticles were loaded with a red light-activated dye Nile Blue with pronounced solvatochromic properties for diagnostic applications and with a green light-activated photodynamic sensitizer Rose Bengal for photodynamic therapy. Targeted delivery was ensured by the non-covalent decoration of nanoparticles with anti-HER2 antibodies, which can be readily adapted for large-scale biotechnological production. *In vitro* and *in vivo* studies proved the effectiveness of red light-mediated HER2-specific imaging and green light-induced cytotoxicity of anti-HER2 PLGA. Interestingly, these particles fluoresced only after cellular internalization, which minimizes background signals inside the organism, thus facilitating real-time diagnostics. The particles allowed efficient and selective visualization of the HER2-positive primary tumor node and metastases spread and led to complete remission in BALB/c Nu/Nu mice with the HER2-positive xenografts after a single session of photodynamic therapy.

## 1. Introduction

The development of multimodal antitumor agents is at the forefront of modern biomedicine ^1,2^. Traditional methods of both diagnostics and therapy lead to a wide range of side effects that crucially affect a significantly weakened organism. Namely, chemotherapy with doxorubicin leads to high cardio- and hepatotoxicity, various surgical methods break the integrity of blood vessels, increasing the rate of metastasis, and various methods of radiation therapy lead to irreversible changes in the bone marrow ^3,4^. The development of safe and effective methods of systemic scanning and targeted therapy on the organism with oncological processes is one of the greatest challenges facing modern natural sciences ^5,6^.

These challenges led to the development of a relatively new direction in biomedicine, theranostics. Theranostics involves the combination of diagnostic and therapeutic tools on a single platform that allows both visualizing the target tissue and, if necessary, performing the treatment and therapy monitoring, thus implementing the concept of a “magic bullet”, formulated by the founder of chemotherapy, Paul Ehrlich ^7,8^. The most suitable platform for developing theranostic agents are biocompatible and biodegradable nanoparticles (polymer and protein nanoparticles, and/or liposomes), which, due to their structure, can serve as carriers of both diagnostic and therapeutic modalities ^9^. At the same time, decorating the surface of nanoparticles with targeting molecules makes it possible to implement the concept of targeted delivery of therapeutic and/or diagnostic compounds directly to the tissue of interest ^10^. For example, targeted therapy for HER2-overexpressing cancers remains one of the most serious problems of modern molecular oncology ^11^. Nanoparticles based on a copolymer of lactic and glycolic acids, poly(lactic-co-glycolic acid) or PLGA, are widely used in the clinic; more than 20 PLGA-based medications have already been approved by the FDA for human administration ^12^. Such particles are used as carriers of therapeutic compounds for the treatment of prostate and breast cancer (Suprefact Depot, Lupron Depot, Trelstar, Eligard, Decapeptyl), neurologic disorders (Risperdal Consta), opioid dependence treatment (Vivitrol, Sublocade) and other diseases.

However, to date, there are no *targeted* PLGA nanoparticles for clinical use, which is caused by the insufficient effectiveness of existing formulations, low tumor accumulation, problems in the transition from animal studies to clinical use, the complexity of chemical conjugation with targeting molecules on an industrial scale, low batch-to-batch reproducibility, and other issues ^13–18^.

The described problems stimulate fundamental research in the field of designing targeted theranostic agents, and different PLGA-based nanoformulations have been previously developed for imaging and therapy of HER2-positive cancer in mice *via* chemotherapy and photothermal therapy. Namely, chemotherapeutic HER2-targeted PLGA nanoparticles loaded with doxorubicin ^19^ and tamoxifen ^20^ were developed. Moreover, several HER2-targeted PLGA-based nanoplatforms combining diagnostic and therapeutic functions were designed for *in vivo* applications. For example, a conjugate of perfluoropentane-loaded PLGA nanoparticles and trastuzumab was developed for ultrasound and magnetic resonance imaging and photothermal therapy of HER2-positive xenografts ^21^, PLGA-based nanoparticles were developed for photoacoustic and magnetic resonance imaging and gold-nanoparticle induced photothermal therapy ^22^.

However, today photodynamic therapy (PDT) is considered to be more promising and safer than photothermal therapy (PTT), which is confirmed by the presence of a sufficiently large number of PDT methods in clinical practice and only a few cases of PTT ^23,24^. Thus the development of HER2-targeted theranostic PLGA nanoparticles with the light-inducible PDT property is expected to make a significant contribution to personalized medicine. But, to date, there are no such PLGA-based nanoparticles for targeted PDT with proven effectiveness *in vivo*. There are several *in vitro* studies on the development of HER2-targeted nanoparticles for PDT and testing their efficacy on cell culture ^25,26^; and non-targeted structures loaded with a photosensitizer have also been generated ^27^.

Here we developed a theranostic platform based on anti-HER2 PLGA nanoparticles with microenvironment-sensitive fluorescent diagnostic properties and light-inducible cytotoxicity for HER2-overexpressing tumors. The platform allows for visualizing and tracking tumor and metastases localization when exposed to red light irradiation and eliminating HER2-overexpressing tumors *in vivo* when exposed to green laser irradiation. Interestingly, the nanoparticles fluoresce only when internalized into cells, which minimizes background signals inside the organism, thus facilitating real-time diagnostics.

The properties are achieved through the loading of the fluorescent dye Nile Blue (NB) with pronounced solvatochromic properties and the photodynamic sensitizer, Rose Bengal (RB), which produces reactive oxygen species (ROS) upon exposure to green light. Targeting properties were provided by the decoration of the nanoparticles with anti-HER2 antibody trastuzumab, which is incorporated onto the nanoparticle surface during the synthesis without the need for chemical modification with cross-linking reagents such as EDC/NHS, which is of great importance for potential biotechnological production. We show that such particles allow efficient and selective localization of the HER2-positive primary tumor node, as well as metastases, and achieve complete remission in the xenograft tumor model in BALB/c Nu/Nu mice.

## 2. Results

### 2.1. Design of experiment

HER2-specific cancer diagnostics and therapy were implemented according to the following experimental scheme (**Fig. 1**). Co-polymer of lactic and glycolic acids, PLGA, was used as the matrix for the synthesis of theranostic nanoagent. PLGA nanoparticles were synthesized by the “water-oil-water” microemulsion method with the incorporation of water-soluble Rose Bengal into the first “water” phase and water-insoluble Nile Blue (NB, 5-amino-9-(diethylamino)benzo[a]phenoxazin-7-ium perchlorate) into the “oil” phase. The nanoparticles were decorated with the anti-HER2 antibody trastuzumab for HER2-specific cancer cell targeting *in vivo*. When administered systemically into the organism, anti-HER2 PLGA nanoparticles selectively bound to HER2-overexpressing cancer cells in a tumor through the interaction of trastuzumab with HER2 receptors present on cancer cell surfaces.

**Figure 1.**
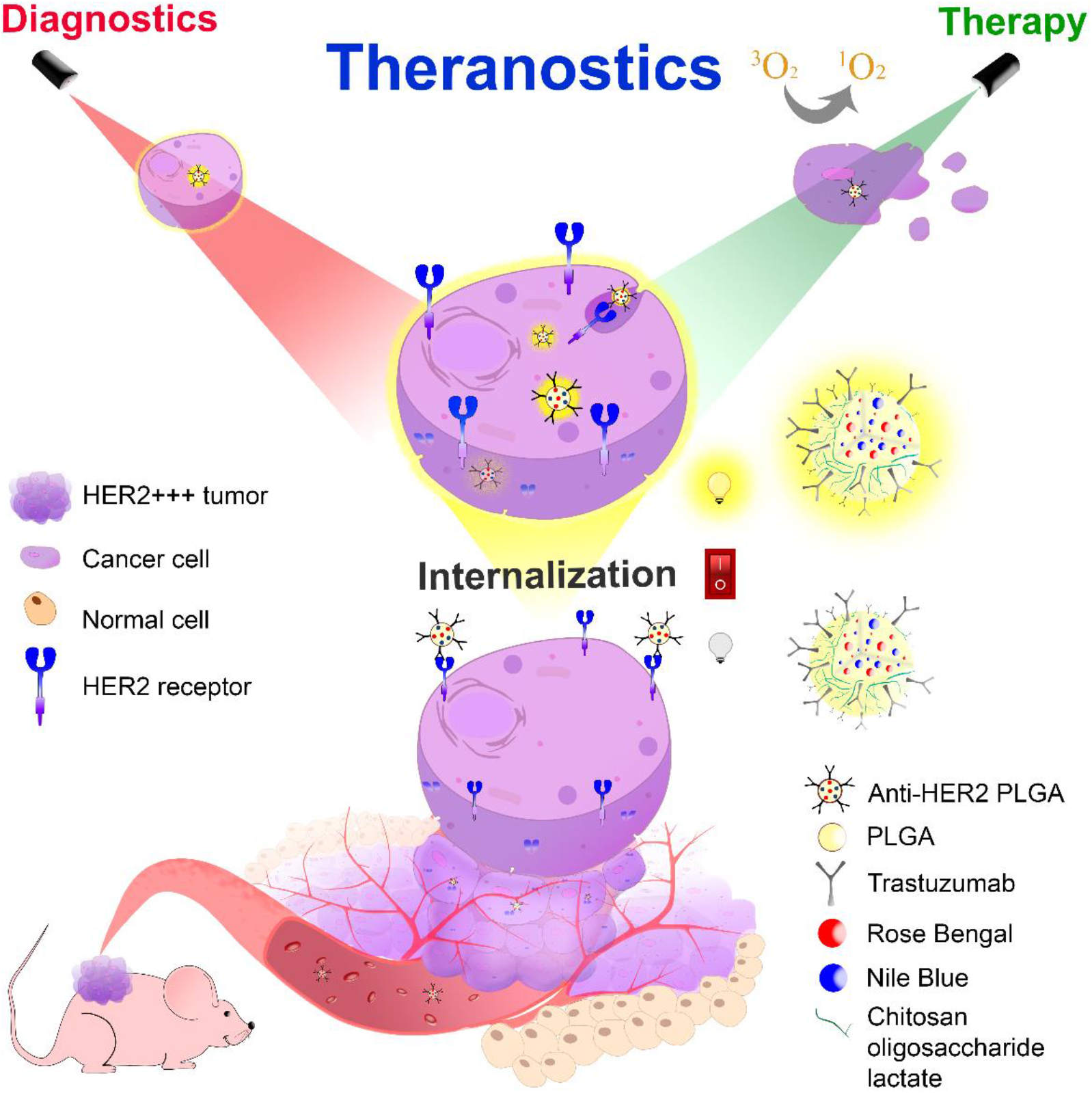
Internalization-responsive nano-PLGA for HER2-positive breast cancer theranostics *in vivo*. Polymer nanoparticles are loaded with the photosensitizer Rose Bengal and fluorescent dye Nile Blue and decorated with the HER2-targeting IgG trastuzumab. The diagnostic dye, Nile Blue, visualizes HER2+ tumor *in vivo* under red light irradiation only after cellular uptake due to solvatochromic properties, while the therapeutic dye Rose Bengal generates reactive oxygen species under green light irradiation. The differential exposure of PLGA to red or green light allows consecutive mapping and elimination of HER2+ cancer cells thus realizing the theranostic concept.

The diagnostic modality of anti-HER2 PLGA nanoparticles was provided through Nile Blue incapsulation followed by red light irradiation for tumor visualization. The nanoparticles were efficiently internalized into cells and after cellular uptake fluoresce upon exposure to red light due to the solvatochromic properties of NB ^28^. NB is a water-insoluble dye that fluoresces only in the lipophilic microenvironment inside cells with weak fluorescence outside cells in water buffer systems (with negligible fluorescence in polar aqueous media outside the cells). This diminished the background fluorescence of the nanoparticles, so tumor visualization did not require their complete elimination from the bloodstream after systemic administration *in vivo*.

The image-guided therapeutic modality was provided through the incorporation of Rose Bengal dye (RB, 4,5,6,7-tetrachloro-2’,4’,5’,7’-tetraiodofluorescein) in the first water phase during the nanoparticle fabrication. RB is a xanthene photosensitizer that produces reactive oxygen species upon exposure to green light. It shows weak fluorescence upon exposure to both green and red light ^29^, does not interfere with the fluorescence of NB, and does not generate ROS under red light thus being an ideal PDT agent that can be activated after NB-mediated bioimaging. Thus, the diagnostic and therapeutic modalities are combined within the single nanoparticle fabricated for differential activation with red or green light irradiation thus providing image-guide HER2-specific PDT treatment.

### 2.2. Synthesis and characterization of anti-HER2 PLGA nanoparticles

Polymer nanoparticles were synthesized by the double emulsion method as schematically illustrated in **Fig. 2a**. The first emulsion was obtained by sonication of the mixture of PLGA and Nile Blue (NB) solution in chloroform with the solution of Rose Bengal (RB) in water. The second emulsion was obtained by sonication of the first emulsion and the mixture of poly(vinyl alcohol) and chitosan oligosaccharide lactate in PBS.

**Figure 2.**
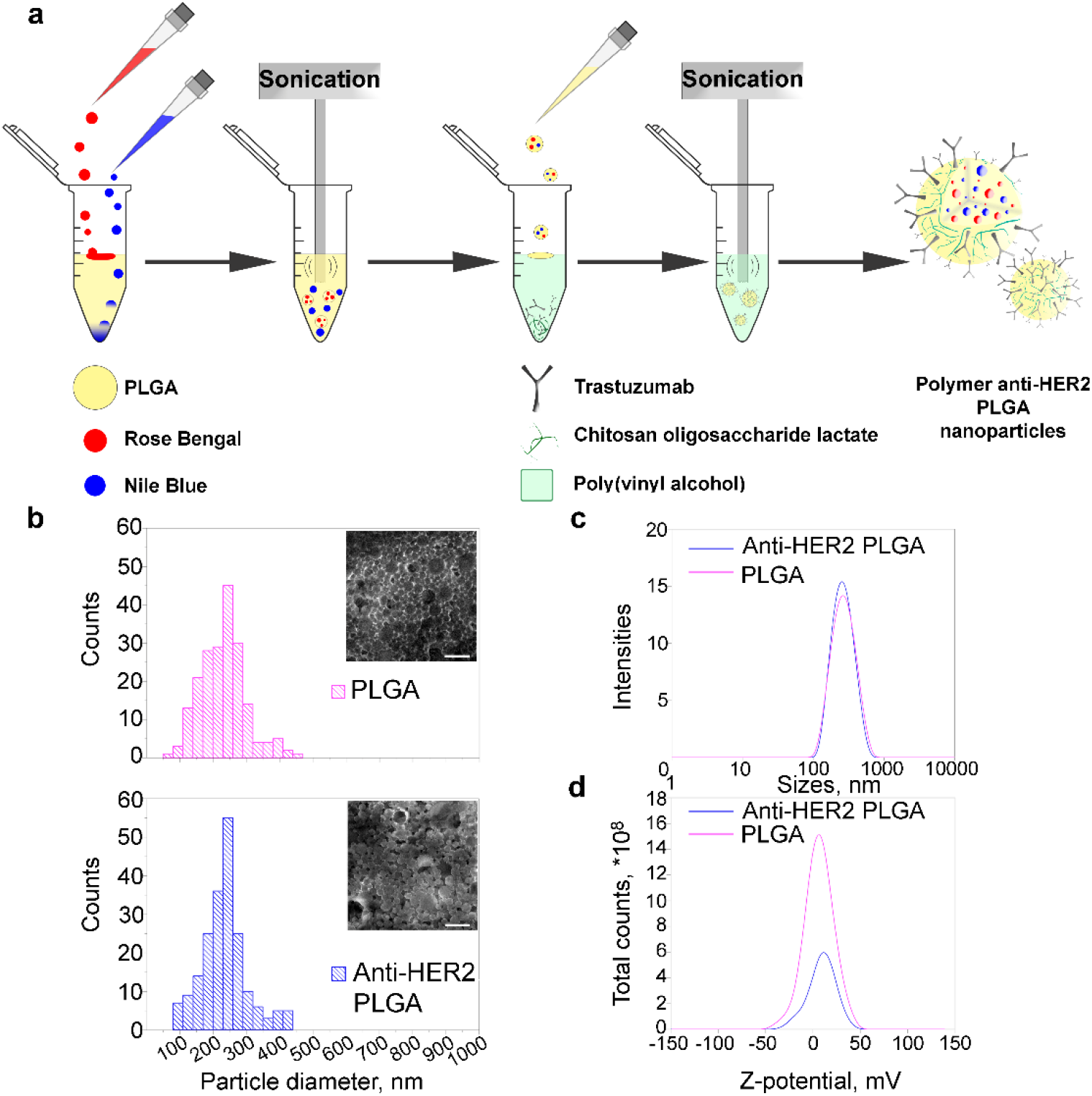
Synthesis and characterization of theranostic anti-HER2 PLGA nanoparticles. (**a**) Schematic illustration of anti-HER2 targeted polymer nanoparticle synthesis *via* “water-oil-water” double emulsion method. Particles are loaded with Nile Blue as a diagnostic modality and with Rose Bengal photosensitizer as a therapeutic modality. (**b**) Scanning electron micrographs and physical size distribution of unmodified PLGA nanoparticles and anti-HER2 PLGA. Scale bar, 1 µm. (**с**) Hydrodynamic size distribution of pristine PLGA particle and anti-HER2 PLGA). (**d**) ζ-potential distribution of unmodified PLGA and anti-HER2 PLGA nanoparticles obtained with electrophoretic light scattering method.

To provide targeting modality, the anti-HER2 antibody, trastuzumab, was added to the second water phase for incorporation into the outer nanoparticle layer likely through its slightly hydrophobic Fc-fragment. The morphology of thus obtained nanoparticles with and without trastuzumab was investigated with scanning electron microscopy (SEM). Data presented in **Fig. 2b** demonstrate that the nanoparticles are monodisperse and spherical with an average diameter of 235 ± 68 nm and 232 ± 67 nm for anti-HER2 PLGA and PLGA, respectively.

The dynamic and electrophoretic light scattering measurements demonstrated the hydrodynamic size of nanoparticles to be equal to 369 ± 166 nm and 323 ± 113 nm for anti-HER2 PLGA and PLGA, respectively (**Fig. 2c)** and ζ-potential –0.66 ± 13.9 m and –3.60 ± 14.8 mV for anti-HER2 PLGA and PLGA, respectively (**Fig. 2d**). The data obtained confirm colloidal stability of as-synthesized nanoparticles. For application *in vivo*, the long-term colloidal stability was confirmed by measuring the hydrodynamic sizes of nanoparticles after 1-month of storage in buffer systems at pH ranging from 2 to 9. To avoid any interference of Nile Blue with the 633 nm laser wavelength of the ZetaSizer NanoZS analyzer, NB-lacking nanoparticles were used (PLGA/RB). The DLS measurements showed the nanoparticles’ stability for a month at pH ranging from 2 to 9; nanoparticles did not form aggregates for at least 35 days under all tested conditions (**Fig. S1**).

### 2.3. Theranostic anti-HER2 PLGA nanoparticles specifically target HER2-positive cells *in vitro*

The targeting efficiency of anti-HER2 theranostic PLGA nanoparticles was confirmed *in vitro* by the specific fluorescent labeling of HER2-overexpressing cancer cells. BT-474 cells with stable expression of luciferase NanoLuc, BT-NanoLuc cells, were used for these tests ^19,30,31^. To verify the HER2 overexpression on the BT-NanoLuc cells’ surface, flow cytometry tests were performed with the addition of the anti-HER2 antibody conjugated with fluorescent FITC dye, trastuzumab-FITC. Chinese hamster ovary (CHO) cells without HER2 (overexpression) were used as a negative control ^32^.

Data presented in **Fig. 3a** confirm the overexpression of HER2 on BT-NanoLuc cells, namely, the median fluorescence intensities (MFI) of cell populations labeled with trastuzumab-FITC were equal to 207 441 and 12 706 for BT-NanoLuc and CHO cells, respectively. The flow cytometry data are supported by the imaging tests: cells were labeled with trastuzumab-FITC and Hoechst 33342 and visualized with confocal laser scanning microscopy (**Fig. 3b**).

**Figure 3.**
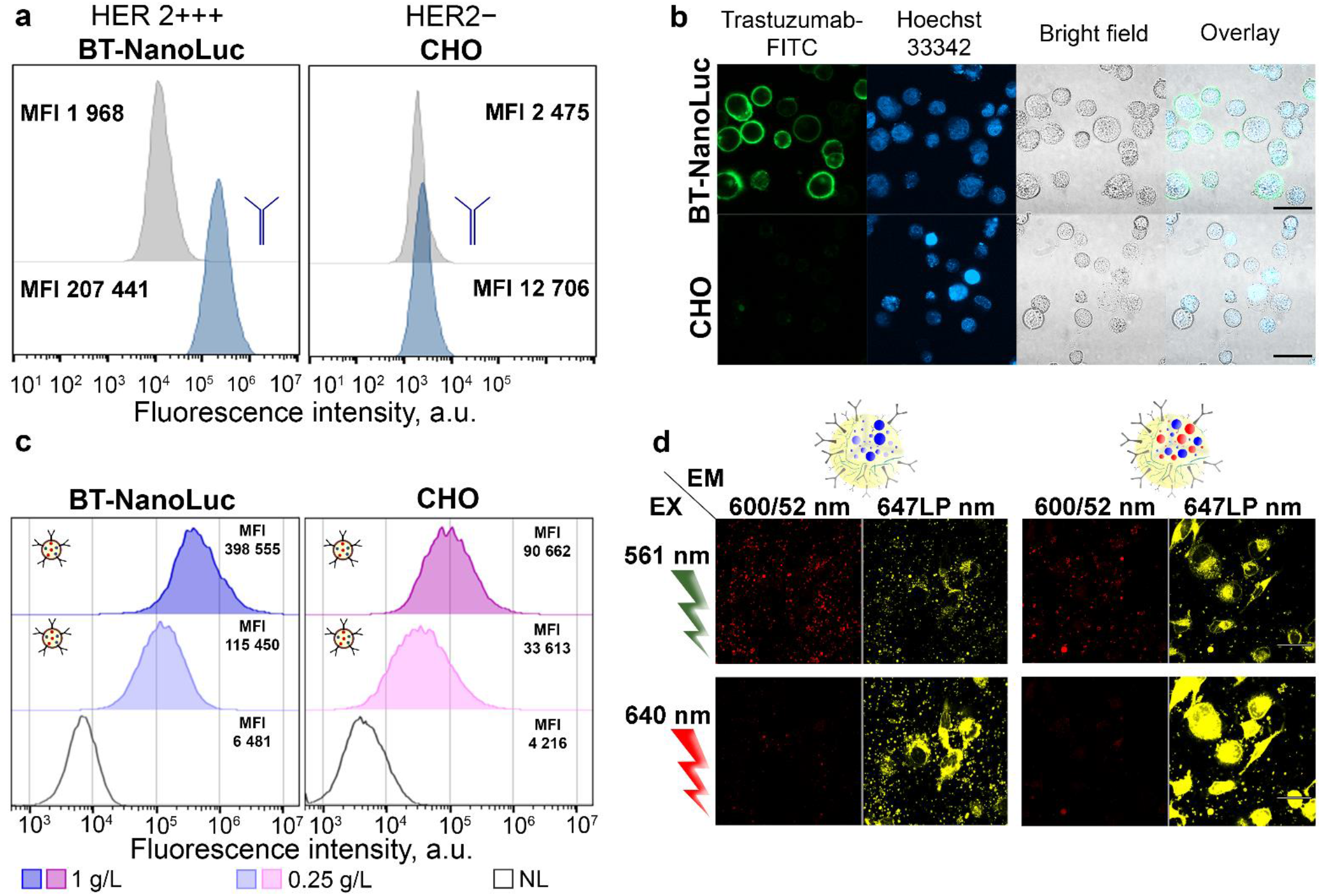
Cancer cell targeting with anti-HER2 theranostic PLGA nanoparticles *in vitro*. (**a**) Flow cytometry analysis of HER2 expression on the surface of BT-NanoLuc and CHO cells. Blue histograms – cells labeled with trastuzumab-FITC, grey histograms – autofluorescence. Data were obtained in the FL1 channel (λ_ex_ = 488 nm, λ_em_ = 525/20 nm). (**b**) Confocal microscopy images of BT-NanoLuc and CHO cells in the presence of trastuzumab-FITC (λ_ex_ = 488 nm, λ_em_ = 492-500 nm) and a nuclear staining dye Hoechst 33342 (λ_ex_ = 405 nm, λ_em_ = 410-520 nm). Scale bars, 50 µm. (**c**) Flow cytometry test on a targeting efficiency of anti-HER2 PLGA nanoparticles. Blue and pink histograms: cells labeled with anti-HER2 PLGA in different concentrations, black open histograms – cells’ autofluorescence. Flow cytometry data were acquired in the FL4 channel (λ_ex_ = 640 nm, λ_em_ = 675/25 nm). (**d**) Cellular uptake-responsive fluorescence of anti-HER2 PLGA nanoparticles due to Nile Blue loading. Confocal microscopy of BT-NanoLuc cells after incubation with anti-HER2 PLGA nanoparticles containing fluorescent dyes: Nile Blue (left), and Nile Blue-Rose Bengal (right) without any washing steps from non-bound nanoparticles. Scale bars, 50 µm.

Next, we evaluated the targeting ability of anti-HER2 PLGA using the flow cytometry test. Data presented in **Fig. 3c** (and **Fig. S2** with the wide nanoparticle concentration range) confirm the specific interactions of targeted anti-HER2 PLGA nanoparticles with HER2-overexpressing cells: MFI of BT-NanoLuc cells labeled with anti-HER2 PLGA is *ca*. 4.5 higher than that for CHO cells.

### 2.4. Fluorescence of anti-HER2 PLGA nanoparticles is cell uptake-responsive

The lipophilic nature of Nile Blue and its highly solvatochromic properties, meaning that the fluorescence emission is highly dependent on the solvent used, especially its polarity, make this dye an excellent candidate for nanoparticle imaging only after cellular uptake. This fact was confirmed using both cell-free and *in vitro* tests.

The imaging of HER2-overexpressing cells with anti-HER2 PLGA nanoparticles was performed. Cells were seeded at 96-well plates and anti-HER2 PLGA nanoparticles with different dye loadings – with NB only with NB/RB were added to cells in equal concentrations. Cells were incubated with nanoparticles for 1 h and then visualized with confocal laser scanning microscopy without any washing steps under the following conditions: λ_ex_ = 561 and 640 nm, respectively, and λ_em_ = 600/52 nm and λ_em_ = 647LP nm (**Fig. 3d**).

The data presented in **Fig. 3d** indicate that upon excitation with a green laser 561 nm, the fluorescence of nanoparticles is observed only outside the cells in the channel λ_em_ = 600/52 nm (pseudo-colored with red) and fluorescence only inside the cells in the 647 LP channel (pseudo-colored with yellow) both for PLGA/NB and PLGA/NB/RB nanoparticles. At the same time, upon λ_ex_ = 641 nm with red laser, the fluorescence of nanoparticles is observed only in the 647 LP channel and corresponds only to nanoparticles internalized inside cells. Thus, by selecting the proper wavelengths of fluorescence excitation and emission, we showed that only nanoparticles internalized into the cell can be detected.

It is also important to note that the presence of Rose Bengal inside nanoparticles enhances this effect. The fluorescence of nanoparticles inside the polar microenvironment (here, cell compartments) was significantly higher for particles loaded with both NB and RB dyes in comparison with nanoparticles loaded with RB only (however, the nanoparticles loaded with RB only did not fluoresce in both channels, see **Fig. S4**). This phenomenon is most likely due to the FRET effect between NB and RB inside the nanoparticles (see figure **Fig. S5**).

### 2.5. Anti-HER2 PLGA nanoparticles provide light-induced tunable cytotoxicity to HER2-overexpressing cancer cells *in vitro* through ROS generation

Along with an effective imaging dye, NB, anti-HER2 PLGA nanoparticles were loaded with RB dye, which is a photosensitizer and is used as an antibacterial agent due to the ROS generation, as well as in studies for melanoma treatment. To find the most effective RB concentration during the nanoparticle synthesis and prevent issues associated with dye aggregation and stacking inside the nanoparticle architecture, the ROS generation efficacy study was performed. Particles were synthesized with dyes at a concentration of 1 g/L NB and a serial 2-fold dilution of RB within the 0.2 ÷ 3 g/L range, as well as particles synthesized with only one dye, NB or RB.

For the preliminary assessment of the therapeutic properties of the targeted particles, the ROS generation was measured when the targeted particles were exposed to green laser light, λ_ex_ = 532 nm. Due to the known properties of Nile Blue to act as a photosensitizer under the 808 nm laser irradiation ^33^, λ_ex_ = 808 nm and the combination of λ_ex_ = 532 and 808 nm were used for nanoparticle irradiation to verify the combination effect of green and NIR irradiation for photosensitization. The total reactive oxygen species assay kit, 520 nm (Thermo) was used to assess the efficacy of ROS generation *via* the fluorescence intensity measurement. According to the data obtained (**Fig. 4a**), the higher the RB concentration, the higher ROS production under λ_ex_ = 532 nm irradiation. Interestingly, the addition of NB into nanoparticles does not affect the ROS generation under the green light irradiation and slightly decreases ROS when the combination of green&NIR light is used (**Fig. 4a**).

**Figure 4.**
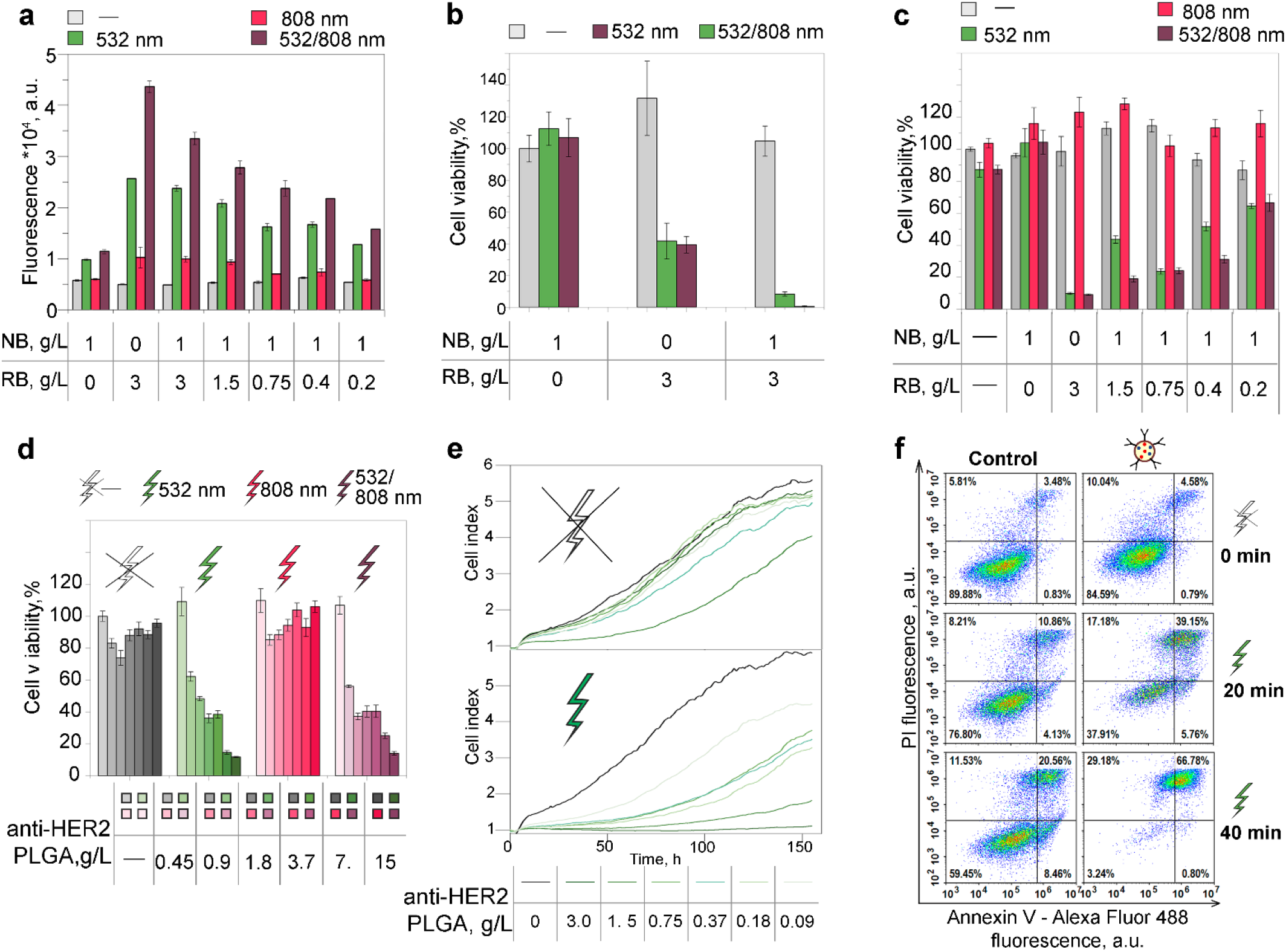
Light-induced cytotoxicity of anti-HER2 PLGA nanoparticles to HER2-overexpressing cancer cells *in vitro* through ROS generation. (**а**) ROS generation by targeted PLGA nanoparticles upon the irradiation with lasers: λ_ex_ = 532 nm, 808 nm, or 532/808 nm, dependence on RB and NB loading into PLGA nanoparticle during synthesis. ROS was measured with a total reactive oxygen species assay kit, 520 nm (Thermo). (**b**) BT-NanoLuc (HER2-overexpressing cells) viability was assessed with the MTT test. Cells were labeled with different nanoparticle formulations and irradiated with lasers: λ_ex_ = 532 nm, 808 nm or 532/808 nm for 10 min. Unlabeled and non-irradiated cells were used as control. (**c**) BT-NanoLuc viability assay: the influence of RB concentration during the synthesis of PLGA on the nanoparticle cytotoxicity was studied. (**d**) BT-NanoLuc viability assay: the concentration-dependent cytotoxicity of anti-HER2 PLGA nanoparticles loaded with NB (1 g/L) and RB (3 g/L) was studied under the irradiation with lasers: λ_ex_ = 532 nm, 808 nm, or 532/808 nm for 10 min. (**e**) Real-time cell analysis of BT-NanoLuc cell growth rate after the incubation with anti-HER2 PLGA nanoparticles and green light irradiation: the cell index dependence on the time of cultivation. (**f**) Сell death mechanism study. Annexin V/PI flow cytometry analysis of cells treated with anti-HER2 PLGA and irradiated with a green light for 0 – 40 min. Flow cytometry dot plots indicate the intensity of PI (a marker of late apoptosis/necrosis) fluorescence on Annexin V fluorescence (a marker of apoptosis). The viable cells, early apoptotic and late apoptotic/necrotic cells are represented in the bottom left quadrant (Annexin V^−^/PI^−^), bottom right (Annexin V^+^/PI^−^), and upper right (Annexin V^+^/PI^+^) quadrants, respectively.

Next, the viability of HER2-overexpressing cells labeled with anti-HER2 PLGA nanoparticles was measured. To assess the percentage of viable cells when exposed to targeted nanoparticles and external light irradiation, a series of MTT tests and colony formation assays were performed (**Fig. 4b-d, Fig. S5**). To evaluate the cytotoxic effects, the cytotoxicity of targeted nanoparticles containing one (NB or RB) or both dyes (NB/RB) was compared. BT-NanoLuc cells were incubated with nanoparticles and irradiated with green or green/NIR lasers. Cells incubated with nanoparticles without irradiation were used as the negative control. Data presented in **Fig. 4b** demonstrate that in contrast to the cell-free ROS study, particles loaded with both dyes exhibit a stronger cytotoxic effect in comparison with particles with RB only under the irradiation with both green and green/NIR light (**Fig. 4b**). This is probably due to the additional cytotoxic effects of the NB dye when exposed to a NIR light that is not associated with the release of reactive oxygen species; no enhanced cytotoxic effect was observed when RB-bearing nanoparticles were exposed to the NIR light, which confirms the effect of NIR light only in the presence of NB (**Fig. 4b**).

Since RB cytotoxicity is not entirely correlated with *in vitro* ROS production, the effect of RB loading concentration on light-induced cytotoxicity was examined. The data presented in **Fig. 4c** confirm that the concentration of 3 g/L during the synthesis is still optimal, which was used in further experiments, namely in concentration- and time-dependent studies. BT-NanoLuc cells were incubated with anti-HER2 PLGA nanoparticles at different concentrations (**Fig. 4d**) and for different incubation times (**Fig. S6**), washed from non-bound nanoparticles, and irradiated with the external laser light. The higher concentration led to higher light-induced cytotoxicity (**Fig. 4d**). Moreover, the anti-HER2 PLGA-induced cytotoxicity increased significantly when the nanoparticles enter the cell – the cytotoxicity is significantly higher after the prolonged incubation of nanoparticles with cells at +37 ºC, and the cytotoxicity can be slightly increased using the combination of green and NIR light irradiation (**Fig. S6**).

### 2.6. Light irradiation of cells labeled with anti-HER2 PLGA causes cell death *via* apoptosis

To study the mechanism of HER2-positive cell death after light irradiation, we first examined the dynamics of cell growth. The growth rate of HER2-positive cells was investigated using a real-time cell analysis (RTCA) E16 xCelligence system (**Fig. 4e, Fig. S7**). BT-NanoLuc cells were incubated with anti-HER2 PLGA nanoparticles, and irradiated with a green laser. Irradiated and non-irradiated cells with nanoparticles were seeded into E16 plates and allowed to grow for 150 h (6.2 days). Data presented in **Fig. 4e** demonstrate that under irradiation, the dynamics of cell growth depend significantly on the nanoparticle concentration. However, in the absence of irradiation, the cell growth is slightly affected by targeted nanoparticles, the only significant decrease is observed for the highest concentration used. The data presented in **Fig. S7** demonstrate that the cell growth dynamic reduction after the irradiation is related mainly to RB loading into the nanoparticle and slightly affected by NB loading (**Fig. S7**).

The obtained data indicate that cell death is caused most probably by the quite slow mechanism of programmed cell death (apoptosis) and not by immediate death due to injury (necrosis). To determine the mechanism of cell death, the Annexin V-Alexa Fluor 488/propidium iodide flow cytometry test was used. BT-NanoLuc cells were incubated with anti-HER2 PLGA nanoparticles at various concentrations, irradiated with a green laser for 0 to 40 min, and seeded into 6-well plates. Cells were cultured for 12 h, and then stained with Annexin V-Alexa Fluor 488/propidium iodide and analyzed with flow cytometry (**Fig. 4f, Fig. S8**). The data presented in **Fig. 4f** demonstrate the presence of apoptotic (right bottom quadrant) and late apoptotic/necrotic (right top quadrant of flow cytometry dot plot) cells depending on the nanoparticle concentration and irradiation time thus confirming the cell death route *via* apoptosis.

### 2.7. Fluorescent bioimaging of HER2-overexpressing tumors with anti-HER2 PLGA nanoparticles *in vivo*

The diagnostic properties of anti-HER2 PLGA nanoparticles were studied by their ability to visualize the localization of HER2-overexpressing tumors *in vivo*. BALB/c Nu/Nu mice underwent subcutaneous transplantation with BT-NanoLuc cells that possess HER2 overexpression. The stable expression of luciferase NanoLuc allows tracking the tumor growth progress and metastasis spreading non-invasively *in vivo* through the intravenous injection of h-coelenterazine – specific substrate of NanoLuc luciferase. When the tumor volume reaches 60 mm^3^, mice were injected with h-coelenterazine and the bioluminescence of the primary tumor on the right flank and metastases on the opposite flank was registered with the IVIS Spectrum CT bioimaging device (**Fig. 5a**). Next, anti-HER2 PLGA nanoparticles were injected at different doses and the fluorescence (λ_ex_ = 640 nm, λ_em_ = 680 nm) was registered after 1h, 2h, 6h, 12h, 18h of injection. Data presented in **Fig. 5a** illustrate the accumulation of anti-HER2 PLGA within the primary tumor area as well as in metastases with significant fluorescence quenching after 2-6 h of injection depending on the injected dose. However, the highest injected dose resulted in stable fluorescence even up to 18 h post-injection. However, the dose of 3 mg was chosen as the optimal one for further tests since it results in stable and informative fluorescence for more than 2 h after injection.

**Figure 5.**
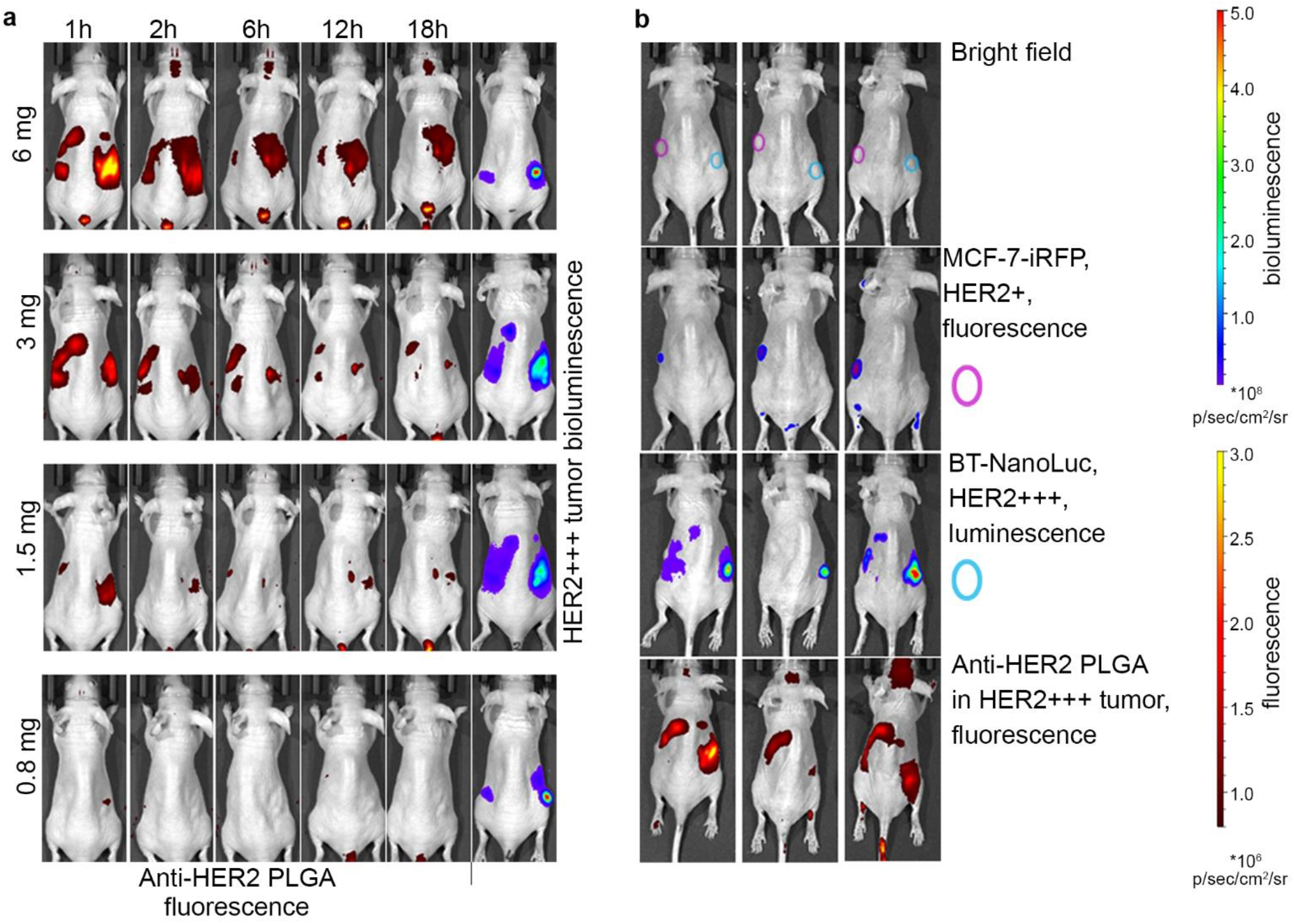
Bioimaging properties of anti-HER2 PLGA nanoparticles *in vivo*. (**a**) BALB/c Nu/Nu mice with s.c. BT-NanoLuc tumors possessing HER2 overexpression were injected with anti-HER2 PLGA and visualized 1, 2, 6, 12, and 18 h post-injection (λ_ex_ = 640 nm, λ_em_ = 680 nm). The primary BT-NanoLuc tumor and metastases are visualized through the i.v. injection of h-coelenterazine and subsequent bioluminescence *in vivo* imaging (right panel). (**b**) Mice with two tumors: 1) MCF7-iRFP tumors (left flank, pink circle, imaged *via* fluorescence detection) and 2) BT-NanoLuc tumors (right flank, blue circle, imaged *via* bioluminescence detection) were injected with anti-HER2 PLGA and 1h post-injection the fluorescence of anti-HER2 PLGA was registered (λ_ex_ = 640 nm, λ_em_ = 680 nm).

Since the HER2 receptor is typically expressed in normal human tissues but in a significantly lower amount compared to breast cancer cells ^19,34,35^, the issue of the selectivity of as-synthesized anti-HER2 PLGA in terms of *in vivo* HER2-binding should be resolved prior to therapy study. To address this question, we used BALB/c Nu/Nu mice with two tumors possessing different HER2 expression. BT-NanoLuc cells with HER2 overexpression were s.c. injected into the right flank, and MCF-7-iRPF fluorescent cells with normal HER2 expression were injected into the left flank. The distribution of BT-NanoLuc cells is registered through the bioluminescence detection, while the localization of MCF-7-iRFP cells is registered through the fluorescence of iRFP protein detection. These mice underwent the injection of anti-HER2 PLGA followed by fluorescent bioimaging (**Fig. 5b**). Data presented in **Fig. 5b** confirm that anti-HER2 PLGA localized predominantly in BT-NanoLuc tumors in the right flank and metastasis on the opposite (left sight) – in inguinal and axillary lymph nodes caused bu BT-NanoLuc tumor with non-significant accumulation in MCF-7-iRFP tumors located between these two lymph nodes.

### 2.8. Anti-HER2 PLGA-based photodynamic image-guided therapy eradicates HER2+ xenogeneic ductal carcinoma *in vivo*

The therapeutic efficacy of anti-HER2 PLGA was assessed *in vivo*. Treatment of xenograft tumors was carried out on day 14 when the tumor volume reached ca. 60 mm^3^ according to the scheme (**Fig. 6a**).

**Figure 6.**
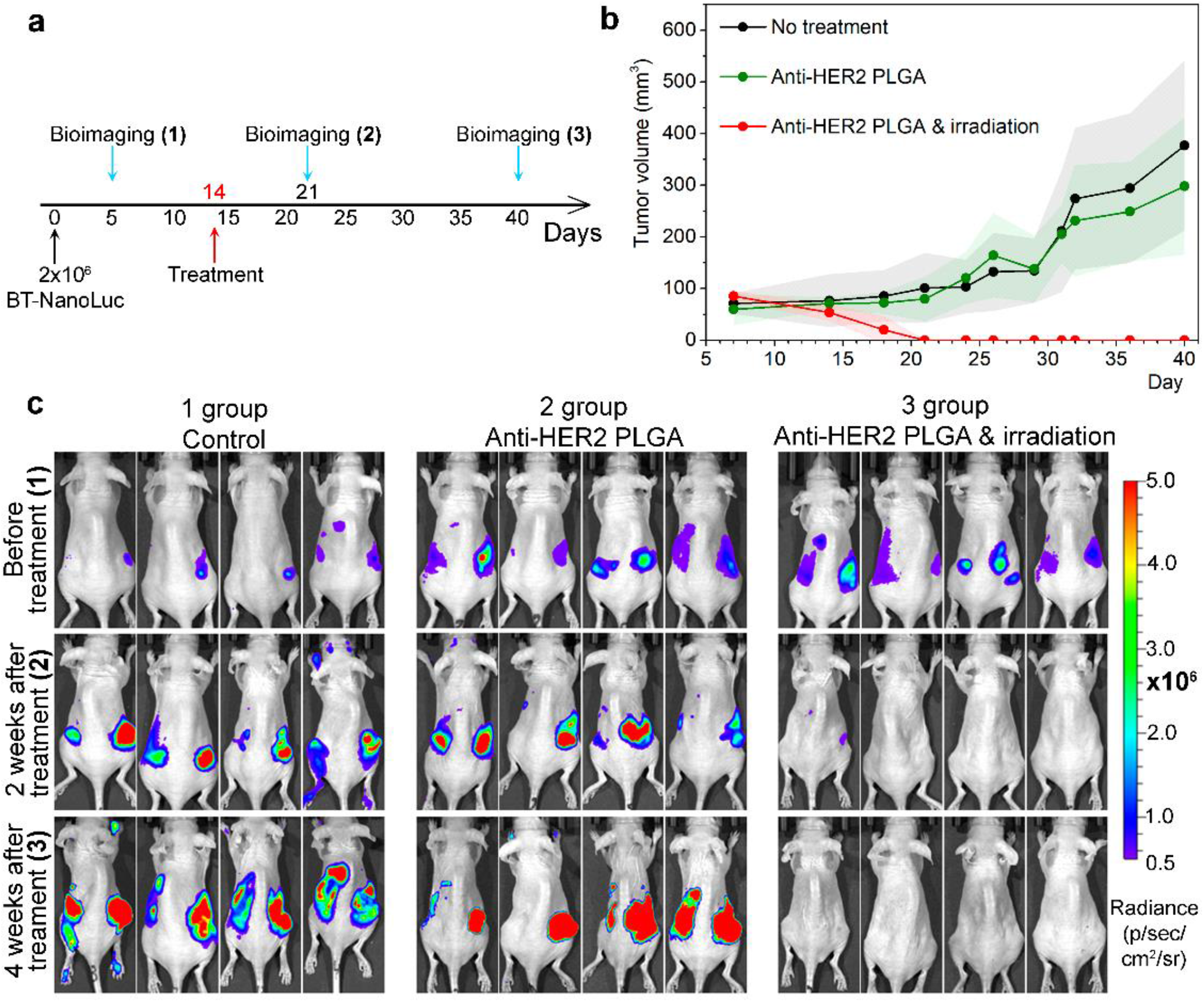
Photodynamic therapy of HER2-positive tumors realized with theranostic anti-HER2 PLGA nanoparticles leading to cancer remission. (**a**) A treatment scheme based on anti-HER2 PLGA. BALB/c Nu/Nu mice were s.c. implanted with 2·10^6^ BT474-NanoLuc cells. After 14 d of tumor inoculation, mice were randomized into groups (n = 4) that received either the injection of anti-HER2 PLGA or a combination of anti-HER2 PLGA with light irradiation. (**b**) Tumor growth dynamics: control group, the group with the single injection anti-HER2 PLGA, and the group with single injection anti-HER2 PLGA and the green light irradiation of primary tumor and metastases. (**c**) Bioimaging of BT474-NanoLuc tumor xenografts: (1) before treatment, (2) and (3) – after treatment. BALB/c Nu/Nu mice were injected with 5 µg of h-coelenterazine, and bioluminescence was recorded with IVIS Spectrum CT.

Mice were randomly divided into three groups (n = 4) and treated with i) PBS, ii) anti-HER2 PLGA, and iii) anti-HER2 PLGA with green light irradiation (**Fig. 6a**). The primary tumors were irradiated for 10 min and areas with metastasis for 3 min. The tumor growth measurements with the caliper (**Fig. 6b**) were accompanied by the bioluminescent *in vivo* imaging (**Fig. 6c**).

*In vivo* bioimaging of experimental animals demonstrate the high therapeutic potential of anti-HER2 PLGA NPs. Tumor growth was fully inhibited by the combination of ani-HER2 PLGA and light irradiation – no luminescence of tumors and metastases was observed by the end of the experiment (40 days after the treatment started).

## 3. Discussion

Theranostic agents are powerful synthetic compounds capable of selective elimination of tumor-cell populations and/or inhibiting their growth and proliferation. The development of a new generation of these agents is critical for the advancement of cancer diagnosis and treatment, as well as for theranostic applications ^36,37^. At the same time, the development and translation into clinical practice of oncotheranostic agents are significantly limited by a number of intractable problems. Such agents are complex multicomponent systems that are not only difficult to synthesize but also difficult to thoroughly characterize, obtaining drug samples with acceptable batch-to-batch reproducibility. In particular, methods of chemical conjugation of nanoparticles with targeting molecules, using, for example, standard cross-linking agents (e.g., EDC/NHS), lead to non-reproducible results of the conjugation reaction due to highly competitive and yet uncontrolled hydrolysis reactions.

These problems are solved by the development of nanoagents that are modified by targeting molecules already during the synthesis process ^38–40^, or by sterically oriented self-assembly of components through various interfaces, such as streptavidin/biotin ^41^, SpyTag/SpyCatcher ^42,43^, barnase/barstar ^44,45^. Considering *in vivo* applications, such approaches, on the one hand, add multifunctionality to drug delivery systems by the possibility of changing self-assembled modules. On the other hand, they significantly affect the size of systems and their immunogenicity, in particular, streptavidin possesses significant immunogenicity ^46^. An alternative solution is the development of fully genetically encoded self-assembling protein nanoparticles with targeted modality which is introduced into the protein’s genetic construct ^35^.

However, considering the loading with therapeutic and diagnostic compounds, protein particles are not as versatile as, for example, liposomes or polymer matrices: upon loading, the tertiary structure of the protein can be disturbed, which affects the efficiency of self-assembly and subsequent possibilities in target recognition. At the same time, the simultaneous loading of hydrophilic and hydrophobic substances into protein particles is a really difficult task while maintaining protein functionality.

To overcome described issues, we developed the method for the synthesis of theranostic polymer nanoparticles decorated with HER2-targeting antibodies without chemical conjugation. For this purpose, a hydrophobic polymer, PLGA, loaded with therapeutic and diagnostic compounds, was used. The surface of PLGA nanoparticles was modified by incorporating the anti-HER2 antibody onto the particle during the microemulsion synthesis – it is most likely, that the Fc fragment of the antibody, due to its inner hydrophobic region, is incorporated into the hydrophobic structure of the PLGA polymer thus covering the surface of nanoparticle with HER2-targeting molecules without the need for chemical conjugation.

PLGA polymer is widely used for extended drug release both in fundamental life science and in clinical practice, and now more than 20 PLGA-based medications are already introduced into the clinic for the treatment of breast and prostate cancer, acromegaly, osteoarthritis, diabetes, growth hormone deficiency treatment, and ocular diseases ^12,47^. PLGA is a biodegradable copolymer that produces lactic and glycolic acids, natural metabolic by-products, upon ester hydrolysis in water ^47,48^. Here we used poly(D,L-lactide-co-glycolide), ester terminated, lactide:glycolide 85:15, Mw 190,000-240,000 polymer. The 85:15 ratio was chosen based on the screening of previously published works. Namely, the ratio of monomers of lactic and glycolic acids in the copolymer affects the rate of hydrolysis, and, accordingly, the time of the release of the loaded substance in the particle. When using polymers, this parameter must be taken into account so that the particle with the substance does not degrade prematurely in the bloodstream before reaching the targeted organ. It has been shown that with a ratio of lactic and glycolic acids in the polymer of 85:15, the release time of the drug in the body increases, compared to 75:25 and 50:50 ratios ^36^. Moreover, the loading of therapeutic substances in a polymer with a ratio of 85:15 is higher than with a 50:50 ratio ^49^.

The “water-oil-water” double microemulsion synthesis method of PLGA nanoparticles proved its versatility for loading different substances – both hydrophobic and hydrophilic ^19,26,50,51^. To incorporate diagnostic modality into nanoparticles, we used the hydrophobic dye Nile Blue, a representative of the benzophenoxazine family, with microenvironment-sensitive fluorescence and high photostability ^52^. The HER2-positive imaging process was guided using the fluorescent properties of Nile Blue under excitation with infrared light. Nile Blue (NB, Nile Blue A, 5-amino-9-(diethylamino)benzo[a]phenoxazin-7-ium perchlorate) is a water-insoluble dye whose excitation and emission enter the NIR-I biological transparency window. NB is being used as a histological stain for *in vitro* and *ex vivo* imaging of lysosomes and lipids ^52^, pH sensorics ^53^, and several *in vivo* studies demonstrated the effective NB accumulation in the tumor after systemic administration ^33,54^. The extensive solvatochromic properties of NB make this dye an excellent candidate for imaging specific cancer cells only after cell internalization. We showed that in aqueous buffer systems the fluorescence of these nanoparticles is very weak, however, after cell internalization, the fluorescence significantly increases in the lipophilic microenvironment inside cells. Considering *in vivo* applications of this property it is worth mentioning that this feature solves the problem of background fluorescence of nanoparticles after systemic administration: when the particles are not completely eliminated from the organism, and the tumor accumulation still did not reach the plateau, the real-time diagnostics is already possible since background fluorescence of circulating particles is not an obstacle for tumor accumulation registration. We demonstrated the applicability of the synthesized targeted nanoparticles for specific visualization of HER2-positive tumors for a long time (up to 18 h) after a single injection into a mouse.

To incorporate therapeutic modality into nanoparticles, we used hydrophilic fluorescent dye, Rose Bengal. Rose Bengal (RB, 4,5,6,7-tetrachloro-2’,4’,5’,7’-tetraiodofluorescein) is a photosensitizer that produces ROS upon exposure to green light ^26,55^. Previously, we experimentally proved that cancer cells can be selectively eliminated when exposed to RB in the PLGA particle and green light irradiation ^26^. RB is a promising diagnostic and therapeutic agent as it is already being used in the treatment of eye diseases ^56,57^ as well as in the treatment of skin cancer ^58,59^. The water-soluble photosensitizer RB fluoresces at 575 nm when exposed to green light at a wavelength of 532 nm. Light with a wavelength of 532 nm is located on the border with the transparency window of biological tissue, however, green light in contrast to red light causes fewer side effects under prolonged exposure ^60,61^. Here, we showed that the treatment with targeted anti-HER2 PLGA nanoparticles loaded with RB and NB and irradiation with a green laser at a wavelength of 532 nm (100 mW), mice with xenograft HER2-positive tumors were completely cured. These data demonstrate the safe and effective application of green light in combination with anti-HER2 PLGA in anticancer PDT since no necrotizing of both tumor and healthy tissue was observed during the treatment.

The obtained nanoparticles were shown to be effective theranostic agents that provide both HER2-specific bioimaging and light-inducible photodynamic therapy *in vivo*. The efficacy *in vitro* and *in vivo* of anti-HER2 PLGA nanoparticles was thoroughly tested, and 100% inhibition of HER2-positive tumor growth in all mice from the experimental group was demonstrated, thus confirming the great potential of the synthesized nanomedication for personalized medicine.

## 4. Materials and Methods

### 4.1. Materials

Sigma, Germany: poly (lactic-co-glycolic) acid (PLGA) (lactide: glycolide 85:15, MW 190-240 kDa), chitosan oligosaccharide lactate, poly (vinyl alcohol) (Mowiol 4-88, PVA, MW 31 kDa), Rose Bengal, Nile Blue A perchlorate; Helicon, Russia: BSA; Chimmed, Russia: chloroform; Roche, Switzerland: Herceptin (Trastuzumab); PanReac Applichem, Germany: dimethylsulfoxide (DMSO).

### 4.2. Synthesis of anti-HER2 PLGA nanoparticles

The targeted particles were synthesized by the «water-oil-water» double emulsion method. The first emulsion was formed by adding 150 µL of 3 g/L Rose Bengal in Milli-Q water and 10 µL of 1 g/L Nile Blue to 300 µL of 40 g/L PLGA in chloroform. The first emulsion was subjected to ultrasound with 750 W Sonicator Vibra-Cell (Sonics & Materials, Inc, USA) for 1 min with 40 % power and 1 min with 60 % power. The second emulsion was prepared by adding the first emulsion to 3 mL of 3 % PVA in water Milli-Q containing 0.3 g/L chitosan oligosaccharide lactate and 200 μL of 10 g/L trastuzumab. The solution was sonicated for 1 min with 40 % power and 1 min with 60 % power at +4 ºC. The chloroform was then evaporated using the rotator at 20 rpm for 3 h at room temperature. The particles were then washed with PBS, pH 7.4 by triple centrifugation and finally resuspended in 100 µL of PBS, pH 7.4. The final concentration of nanoparticles was determined by drying 70 µL of particles at 60 °C.

### 4.3. Scanning electron microscopy

The morphology of particles was evaluated with scanning electron microscopy at an accelerating voltage with a MAIA3 electron microscope (Tescan, Czech Republic) at an accelerating voltage of 5 kV. The physical size of the targeted particles was determined with ImageJ software to get the particle size distribution.

### 4.4. Dynamic and electrophoretic light scattering measurements

The hydrodynamic size of PLGA nanoparticles was determined using a Zetasizer Nano ZS analyzer (Malvern Instruments Ltd, UK) in Milli-Q water at 25 ºC. Ζ-potential was measured using Zetasizer Nano ZS (Malvern Instruments Ltd, UK) in 10 mM KNO_3_ at 25 °C. Particle stability was measured with a Zetasizer Nano ZS (Malvern Instruments Ltd, UK) through consecutive measurements of the size within one month. The particles were diluted in buffer solutions with pH from 2 to 9 and the hydrodynamic size of the particles was measured. The following buffer systems were used: pH 2 (0.1 M glycine-HCl), pH 3 (0.1 M glycine-HCl), pH 4 (0.15 M citrate-phosphate buffer), pH 5 (0.15 M citrate-phosphate buffer), pH 6 (0.1 M HEPES buffer), pH 7 (0.15 M phosphate buffer), pH 8 (0.1 M TRIS buffer), pH 9 (0.1 M TRIS buffer).

### 4.5. Cell culture

The source of cell lines is the collection of the laboratory of molecular immunology, Institute of Bioorganic Chemistry, Russian Academy of Sciences: human breast cancer cells BT-474 with stable NanoLuc expression, BT-NanoLuc ^19^, MCF-7 with iRFP protein expression, MCF-7-iRFP, Chinese hamster ovary, CHO. BT-NanoLuc cells were cultured in DMEM growth medium (Gibco, USA) supplemented with 10% fetal bovine serum (Capricorn, Germany), 2 mМ L-glutamine (PanEko, Russia), 50 U/mL/50 μg/mL penicillin-streptomycin (PanEko, Russia), 1 µg/mL puromycin (Sigma, Germany) and 10 µg/mL ciprofloxacin (Elfa, India) in culture flasks (Nunc, Denmark) in a CO_2_ incubator (BINDER, Germany) at 37 °C with 5 % CO_2_. MCF-7-iRFP cells were cultured in FluoroBright DMEM growth medium (Gibco, USA) with 10% fetal bovine serum (Capricorn, Germany), 2 mM L-glutamine (PanEko, Russia), 50 U/mL/50 μg/mL penicillin-streptomycin (PanEko, Russia) and 10 µg/mL ciprofloxacin (Elfa, India) in culture flasks (Nunc, Denmark) in a CO_2_ incubator (BINDER, Germany) at 37 °C with 5% CO_2_. Cells were removed from the surface of culture flasks using a 2 mM EDTA solution (PanEko, Russia).

### 4.6. Confocal microscopy

Visualization of HER2-overexpressing cells labeled with targeted nanoparticles was performed on the Leica DMI6000B (Leica Microsystems, Germany) microscope equipped with confocal microscopy upgrade (Thorlabs, USA) at the following conditions. BT-NanoLuc cells were seeded at 96-well flat-bottomed culture plates in 100 µL of DMEM growth medium, incubated overnight under a humidified atmosphere with 5 % CO_2_ at 37 °C and anti-HER2 PLGA targeted particles were added in 100 µL of full culture medium to get a final concentration of nanoparticles 10 µg/mL. Cells were incubated with nanoparticles for 1 h at 37 °C, and washed from unbound particles with subsequent microscopy tests. Cells were imaged with green and red lasers separately and in combination, namely: 561 nm green laser excites Rose Bengal dye; 640 nm red laser excites Nile Blue dye. Emission filters 600/52 nm and 647LP nm were used for visualization for all lasers.

Trastuzumab-FITC binding was visualized using an LSM 980 (Zeiss) confocal microscope. Cells were treated with 1 µg/mL Hoechst33342 and 2 µg/mL trastuzumab-FITC in PBS with 1% BSA for 30 min, washed with PBS and imaged under the following conditions: (λ_ex_ = 488 nm, λ_em_ = 492-500 nm for trastuzumab-FITC detection and λ_ex_ = 405 nm, λ_em_ = 410-520 nm for Hoechst 33342 detection.

### 4.7. Fluorescence spectroscopy of targeted nanoparticles

Targeted nanoparticles loaded with Rose Bengal and Nile Blue dyes were resuspended in 100 µL of PBS at 10 µg/mL, pH 7.4, and/or in 100 µL of DMSO and added to the 96-well plate. The particle fluorescence was measured using an Infinite M100 Pro spectrophotometer (Tecan, Austria) within excitation wavelengths 500-600 nm range.

### 4.8. ROS study

The generation of reactive oxygen species by targeted particles was measured using irradiation laser wavelengths of 532 nm and/or 808 nm. Particles were synthesized with different concentrations of dyes. To perform the ROS generation test, 600 µL of PBS pH 7.4 was added to 1.5 mL tubes, next 20 µL nanoparticles at 30 g/L were added, the sample was divided into 4 tubes, and the Invitrogen reagent (Total Reactive Oxygen Species (ROS) assay kit, 520 nm) was added to these samples according to the manufacturer’s recommendations. Then samples were irradiated with lasers individually and in combination (532 nm, 808 nm, 532&808 nm) for 5 min. Then, the fluorescence of irradiated nanoparticles was measured in 96-well plates with 488 nm excitation and 525 nm emission with a gain of 149 on an Infinite M100 Pro spectrophotometer (Tecan, Austria).

### 4.9. MTT assay

The cytotoxicity of nanoparticles under different conditions was determined using the MTT test.

i. The efficacy of dual dye loading into nanoparticles was determined as follows. 75·10^3^ BT-NanoLuc cells were added to 1.5 mL tubes in 300 µL of 1% fat-free milk in PBS, next 25 µL 30 g/L of particles were added, samples were incubated for 15 min at +4 ºC, and washed from unbound particles by triple centrifugation. Finally, cells were resuspended in 600 μL of 1% BSA and divided into 200 μL samples in 1.5 mL tubes. Cells were irradiated for 1) 532 nm laser, 100 mW (KLM-532-200 Optronic, Russia), 10 min, room temperature; 2) 808 nm laser (KLM-H808-1200-5 Optronic, Russia), 1200 mW, 10 min, 30 °C, 3) the combination of both lasers. Next, cells were seeded into 96-well plates at a concentration of 5000 cells per well in 200 μL of complete DMEM medium and cultured for 48 h. Then the medium was removed, 100 μL of MTT solution at 0.5 g/L was added to each well, and plates were incubated for 1 h at 37 ºC and 5% CO_2_. Next, the MTT solution was removed and 100 µL of DMSO was added. The absorbance was measured at a wavelength of λ = 570 nm (reference λ = 630 nm) using an Infinite M100 Pro plate reader (Tecan, Austria).
ii. The concentration-dependent cytotoxicity of anti-HER2 PLGA/NB/RB was determined similarly by incubating cells with 15, 7.5, 3.75, 1.8, 0.9, 0.45 g/L anti-HER2 PLGA/NB/RB.
iii. The cytotoxicity of nanoparticles loaded with different concentrations of RB was determined similarly by incubating cells with particles synthesized with different RB concentrations (3, 1.5, 0.75, 0.4, 0.2 g/L).
iv. The influence of incubation time on cytotoxicity was determined similarly by irradiation cells 1) immediately after the washing from non-bound particles, 2) after 1 h of incubation at 37 ºC, 3) after 4 h after incubation at 37 ºC.

### 4.10. Colony formation assay

BT-NanoLuc cells were incubated with targeted anti-HER2 PLGA/NB/RB particles in different concentrations (15, 7.5, 3.75, 1.8, 0.9, 0.45 g/L) in 1.5 mL test tubes at a concentration of 75·10^3^ in 100 µL of colorless DMEM medium. Incubation was carried out at +4 ºC on a rotator for 15 min. Cells were washed from unbound particles and resuspended in 100 μL of PBS with 1% BSA. Next, the cells with particles were irradiated for 10 min with 532 nm and/or 808 nm for 10 min. Irradiated BT-NanoLuc cells with targeted particles were added to the wells of a 12-well plate: 10^3^ cells in 1200 μL of complete DMEM medium. The plates were incubated at 37 ºC and 5% CO_2_ for 7 days until colonies of more than 50 cells were formed. After incubation, the medium was removed, and the wells were washed with 600 µL of PBS and 600 μL of 70% ethanol in PBS. Next, 600 μL of 95% ethanol was added, and wells were incubated for 15 min. Next, after 95% ethanol was removed, the wells were washed with tap water and 600 μL of 1% crystal violet was added. Wells were incubated for 30 min at room temperature. The crystal violet was then removed and the wells were washed 10 times with tap water. The plates were scanned (EPSON Perfection 2400 Photo, Indonesia).

### 4.11. Real-time cell survival analysis

Real-time cell survival analysis was performed on the xCELLigence instrument (Agilent, USA). 3·10^3^ of BT-NanoLuc cells were incubated with nanoparticles PLGA/RB, PLGA/NB, PLGA/NB/RB, and PLGA/NB/RB) at various concentrations in 100 µL of PBS for 15 min at +4 ºC. Next, the cells were washed from unbound particles 3 times with 200 µL of PBS and resuspended in 300 µL of phenol red-free complete growth medium.

Next, samples were irradiated for 10 min with a 532 nm laser (100 mW). Then, 150 μL of irradiated and non-irradiated cells with particles were added to each well of the E16 xCELLigence plate, into which 50 μL of a complete phenol red-free growth medium DMEM were previously added. The binding kinetics and cell growth were measured for 150 h at 37 ºC and 5% CO_2_.

### 4.12. Apoptosis test

The Invitrogen reagent kit (Molecular Probes Dead Cell Apoptosis Kit with Annexin V Alexa Fluor 488 & Propidium Iodide, PI) was used to study the mechanism of cell death. 400·10^3^ of BT-NanoLuc cells were incubated with anti-HER2 nanoparticles at 2 concentrations, washed from non-bound particles by centrifugation and resuspended in 400 µL of colorless FluoroBrite DMEM medium (Gibco, USA). Next, 500 µL of a FluoroBrite DMEM medium was added to 6-well plates, and 100 µL of 10·10^4^ cells incubated with nanoparticles were added per well. Next, the cells were irradiated for 0, 1, 3, 10, 20, 40 min with a 532 nm. Cells without the addition of particles and without irradiation were used as controls. The plates were incubated for 12 h at 37 ºC and 5% CO_2_. After incubation, the medium was removed from the plates and the cells were detached with 2 mM EDTA solution. Annexin V and PI reagents were added to the obtained samples according to the manufacturer’s recommendations. The samples were then evaluated on a NovoCyte 3000 VYB cytometer (ACEA Biosciences, USA) in the BL1 channel (excitation laser 488 nm, emission filter 530/30 nm) for Annexin V and the BL3 channel (excitation laser 488 nm, emission filter 615/20 nm) for PI.

### 4.13. Tumor-bearing mice

Female BALB/c Nu/Nu mice (20-25 g) were obtained from the vivarium of the Institute of Bioorganic Chemistry, (Pushchino, Russia). The mice were housed in specific pathogen-free facilities with free access to food and water. All procedures were approved by the IBCh RAS Institutional Animal Care and Use Committee (protocol No. 298/2020, dated 29 May 2020). Experiments were performed using Zoletil (Virbak, France) and Rometar (Bioveta, Czech Republic) anesthesia at a dose of 25/5 mg/kg. Aerrane (Isoflurane, Baxter) inhalation was performed for anesthesia using the RAS-4 Rodent Anesthesia System (Perkin Elmer, Waltham, MA, USA) for experiments with IVIS Spectrum CT living imaging system (Perkin Elmer, Waltham, MA, USA).

Xenograft tumors were obtained by inoculating 2·10^6^ BT-NanoLuc cells with 30% Matrigel (Corning, USA) in 100 µL of complete culture medium subcutaneously into the right flank of BALB/c Nu/Nu mice. 2·10^6^ MCF-7-iRFP cells were s.c. inoculated into the left flank of BALB/c Nu/Nu mice.

### 4.14. In vivo bioimaging

Bioluminescent images of xenograft tumors were obtained through i.p. injection of h-coelenterazine (200 µL, 25 µg/mL in PBS). Next, bioluminescent images were captured with an open filter on the IVIS Spectrum CT system (Perkin Elmer, Waltham, MA, USA) IS1803N7357 iKon camera (Andor, USA) using 10-s exposure time (f/stop = 1, binning = 8, the field of view 13.4 cm × 13.4 cm) and normalized to photons per second per cm^2^ per steradian (p/sec/cm^2^/sr). For nanoparticle localization studies, fluorescent images were obtained respectively under the following conditions: excitation 640 nm, emission 680 nm, exposure = 1s, f/stop = 2, binning = 8, the field of view 13.4 cm × 13.4 cm. MCF-7-iRPF tumor imaging was performed under the following conditions: excitation 605 nm, emission 660 nm, exposure = 1s, f/stop = 2, binning = 8, the field of view 13.4 cm × 13.4 cm.

### 4.15. Therapy study

For the treatment experiments, mice were challenged with cancer cells, then randomly divided into three groups (n = 4 in each). The tumor growth dynamic was assessed with the tumor volume measurement using the caliper and tumor volume calculation with the formula V = a^2^·b/2, where a is the width and b is the length of the tumor. When the tumor volume reached 60 mm^3^, mice received an i.p. injection of nanoparticles in PBS. 1h after the injection, the fluorescent images were obtained and mice were irradiated with the light (laser, 532 nm, 100 mW) according to the obtained images. Namely, primary tumors (10 min) and lymph nodes (3 min) with the localization of targeted nanoparticles were irradiated.

### 4.16. Statistical analysis

All experiments were performed independently three times. Numerical data are shown as mean ± standard deviation. Significant differences were determined using a two-tailed Welch t-test. Statistically significant – P <0.05 (*), 0.01 (**), 0.001 (***).

