## Supplementary material for "Internalization-Responsive Nano-PLGA For Image-Guided Photodynamic Therapy Against HER2-Positive Breast Cancer": Fig.S1-S8

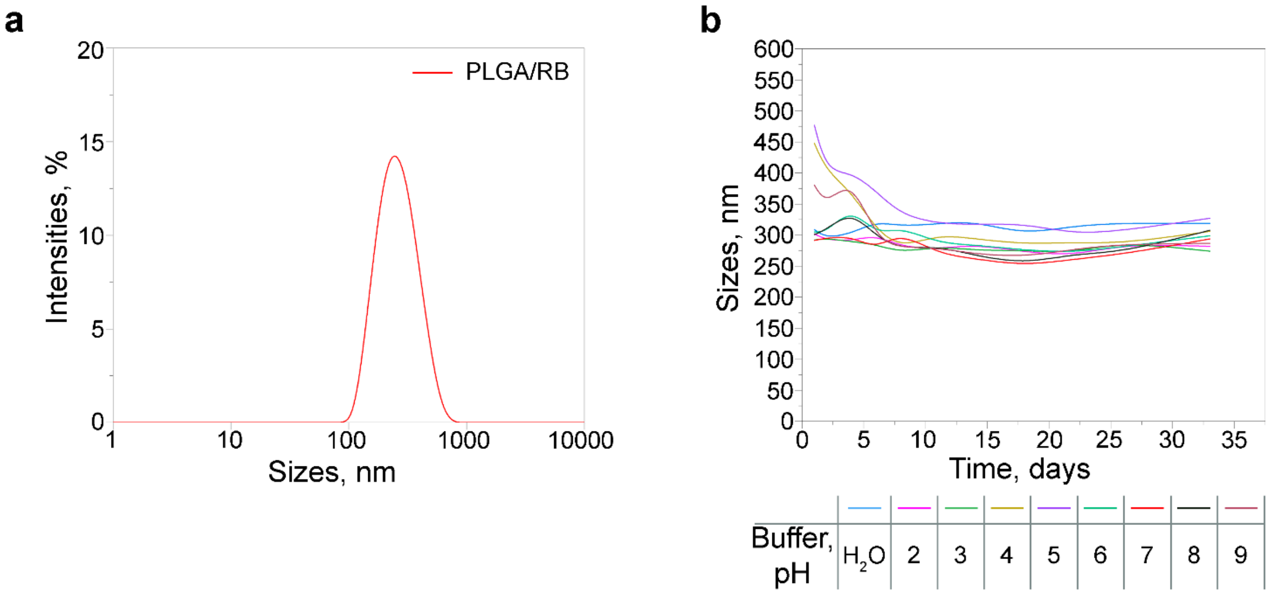


**Figure S1.** PLGA nanoparticle characterization. (a) The hydrodynamic size of PLGA/RB particles. (b) Colloidal stability of PLGA/RB at pH 2-9 measured with DLS.

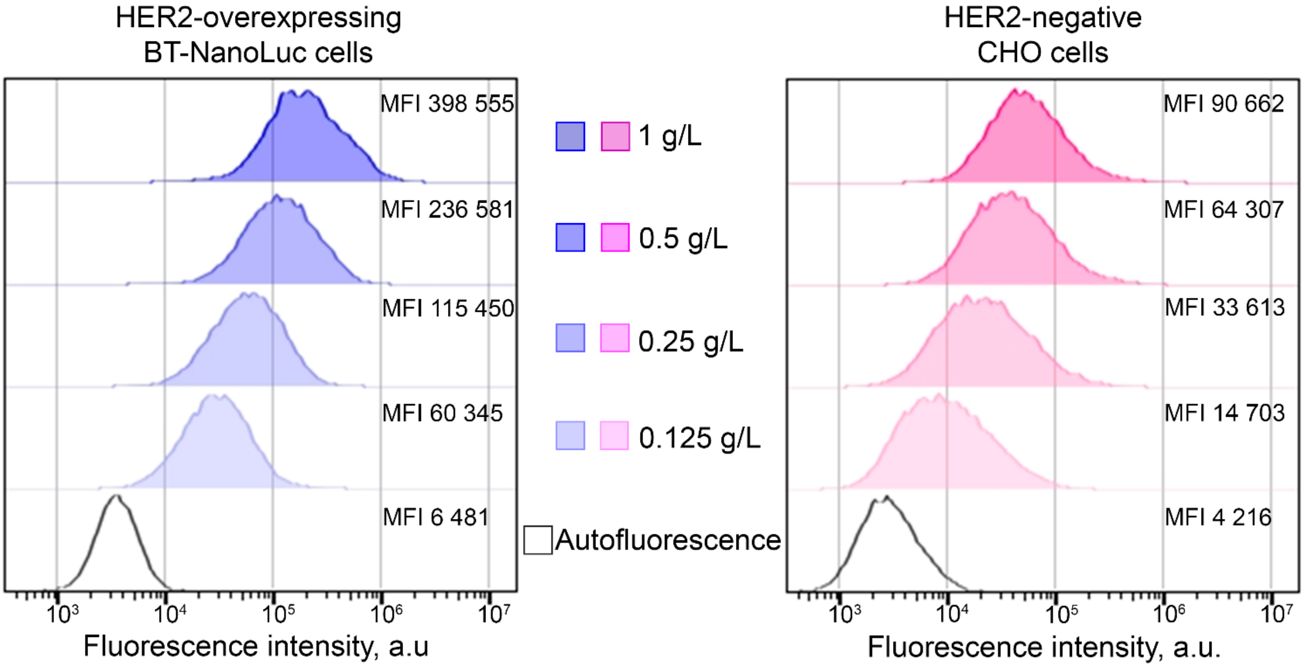


**Figure S2.** Flow cytometry test on anti-HER2 PLGA nanoparticles targeting efficiency: blue and pink histograms: cells labeled with anti-HER2 PLGA in different concentrations, black open histograms – cells' autofluorescence. Flow cytometry data were acquired in the FL4 channel ($\lambda_{ex}$ = 640 nm, $\lambda_{em}$ = 675/25 nm).

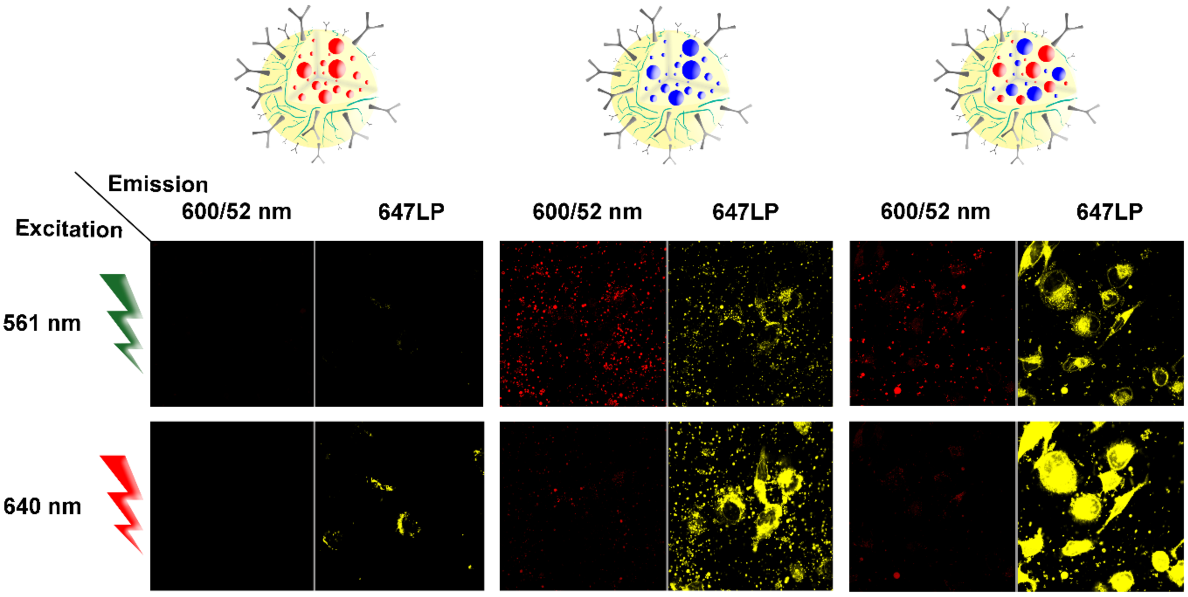


**Figure S3.** Cellular uptake-responsive fluorescence of anti-HER2 PLGA nanoparticles due to Nile Blue loading. Confocal microscopy of BT-NanoLuc cells after incubation with anti-HER2 PLGA nanoparticles containing fluorescent dyes: Rose Bengal (left), Nile Blue (center), and Nile Blue-Rose Bengal (right) without any washing steps from non-bound nanoparticles. Scale bars, 50 µm.

Solvatochromic properties of Nile Blue within nanoparticles were verified by fluorescence spectroscopy measurements. Nanoparticles loaded with Nile Blue – PLGA/NB, Rose Bengal – PLGA/RB, or with both dyes – PLGA/NB/RB were irradiated at $\lambda_{ex}$ = 500-530 nm suitable for Rose Bengal excitation and with $\lambda_{ex}$ = 590-630 nm suitable for Nile Blue excitation in aqueous buffer system (PBS) or polar solvent, dimethyl sulfoxide (DMSO).

Data presented in **Fig. S4** confirm that when nanoparticle is loaded with both dyes, its fluorescence significantly increased compared to the loading with NB or RB alone in polar solvent (see excitation at 630 nm in DMSO) most probably due to FRET effect between RB and NB.

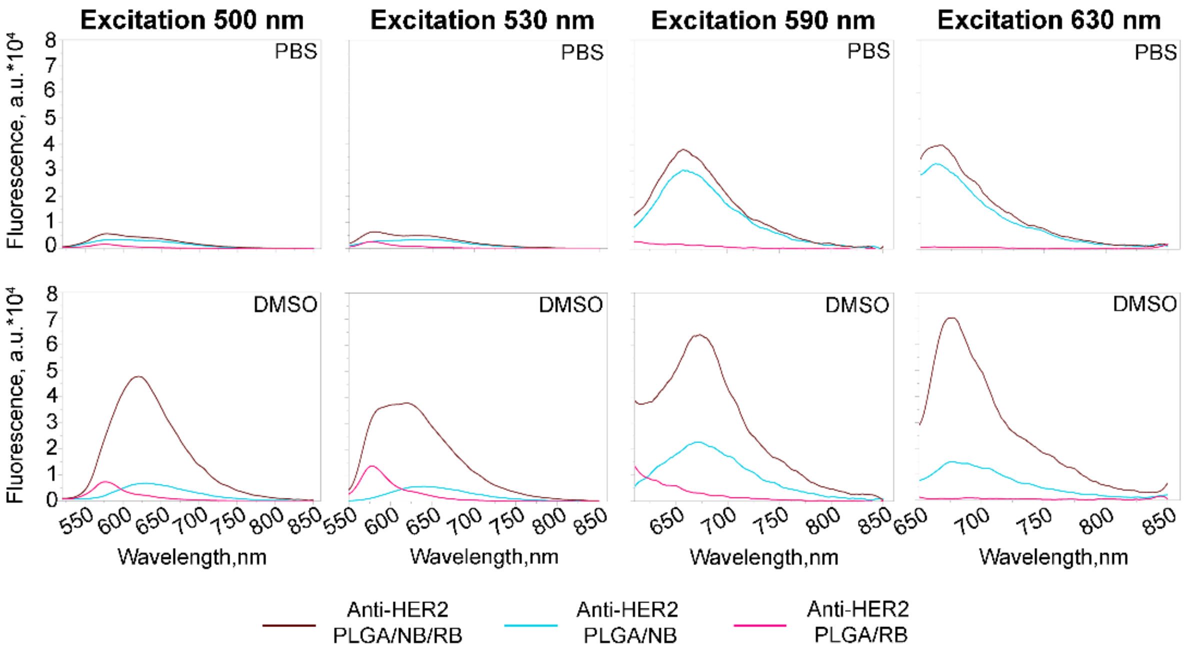


**Figure S4**. Nile Blue-guided solvatochromic nature of anti-HER2 PLGA nanoparticles. Fluorescence emission spectra of targeted anti-HER2 nanoparticles with different dye loadings in PBS and DMSO upon λex = 500 nm to 630 nm were registered.

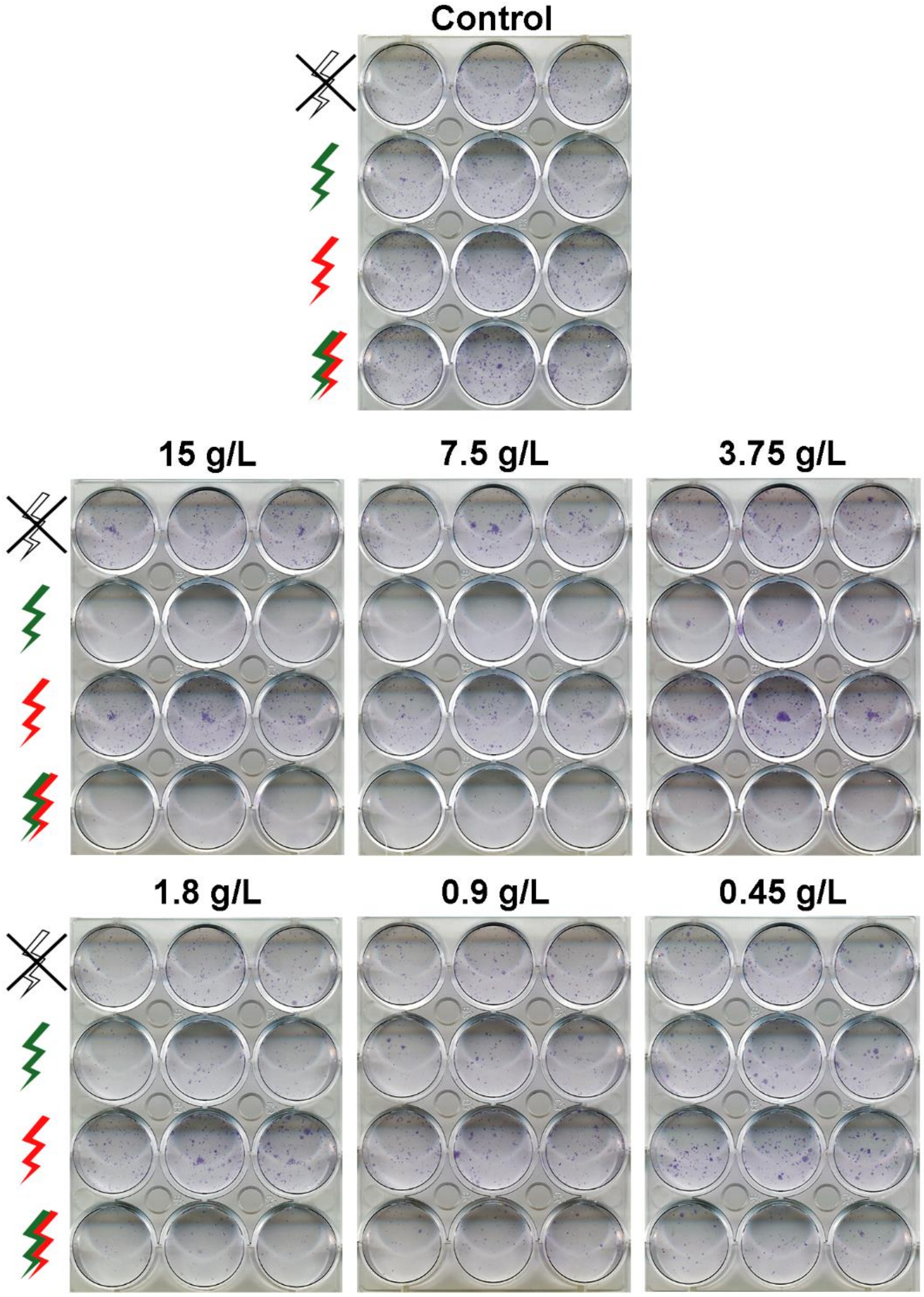


**Figure S5.** Colony formation assay – test on cytotoxicity of anti-HER2 PLGA after irradiation with lasers: $\lambda_{ex}$ = 532 nm, 808 nm or 532/808 nm for 10 min.

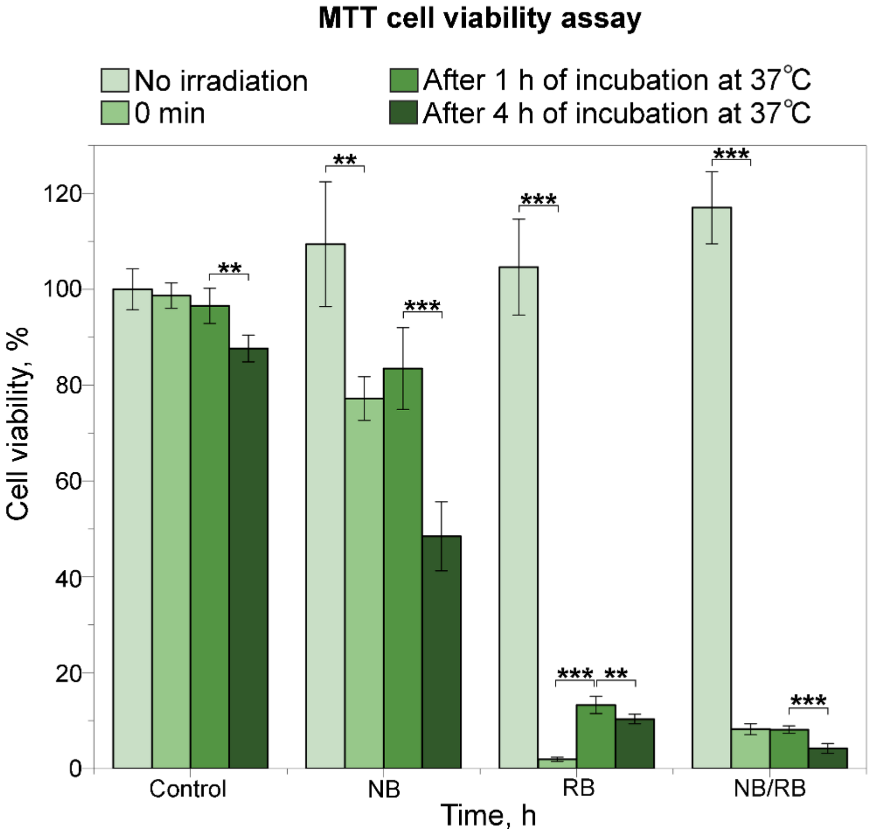


**Figure S6.** Light-induced cytotoxicity of anti-HER2 PLGA nanoparticles to HER2-overexpressing cancer cells *in vitro*. Cells were incubated with targeted nanoparticles with various dye loadings (NB, RB, NB/RB) and irradiated with a green laser for 10 min. Irradiation was carried out immediately (0 min), 1 h, and 4 h after the washing of non-bound particles from cells. Cells without the addition of particles were used as controls. P <0.01 (**), 0.001 (***), Welch's t-test.

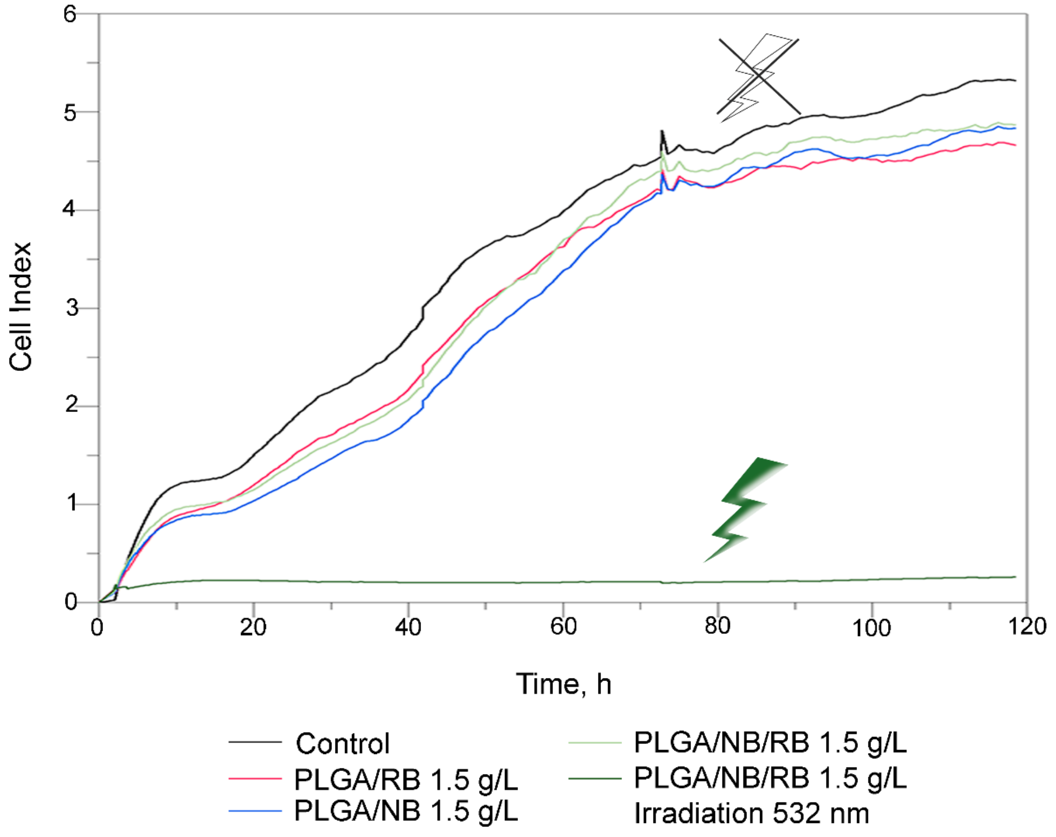


**Figure S7.** Real-time cell analysis of BT-NanoLuc cell growth rate after the incubation with anti-HER2 PLGA nanoparticles and green light irradiation: the cell index dependence on the time of cultivation. Cells were incubated with particles of the same concentration, but with different dye loadings (RB, NB, NB/RB). A sample of cells with PLGA/NB/RB particles was irradiated with a green laser, and the rest of the samples were cultured without irradiation.

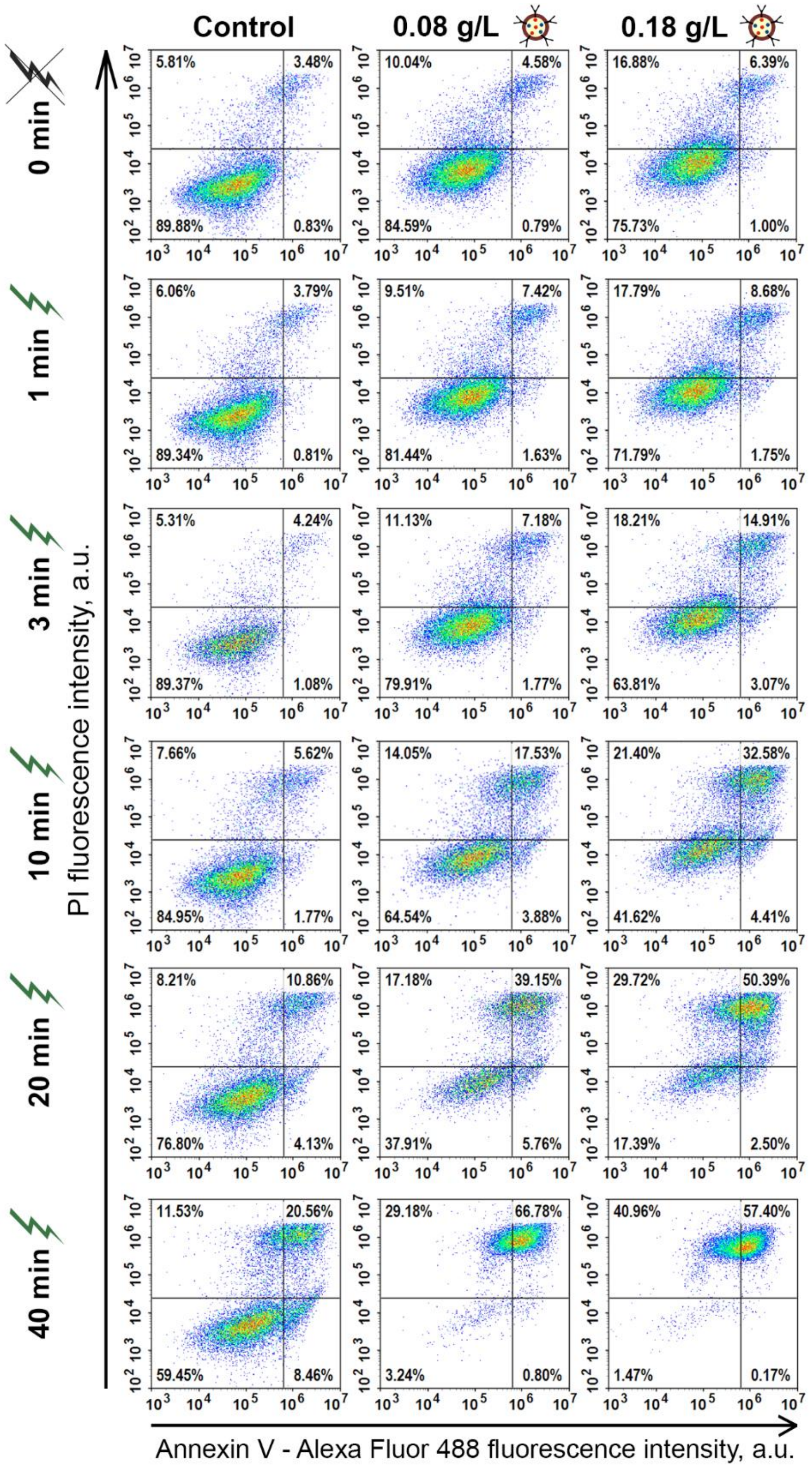


**Figure S8.** Cell death mechanism study. Annexin V/PI flow cytometry analysis of cells treated with anti-HER2 PLGA at different concentrations and irradiated with a green light for 0 – 40 min. Flow cytometry dot plots indicate the intensity of PI (a marker of late apoptosis/necrosis) fluorescence on Annexin V fluorescence (a marker of apoptosis). The viable cells, early apoptotic and late apoptotic/necrotic cells are represented in the bottom left quadrant (Annexin $V^-/PI^-$), bottom right (Annexin $V^+/PI^-$), and upper right (Annexin $V^+/PI^+$) quadrants, respectively.
